# EFFICIENT EXPANSION OF NK-92 CELL LINE USING A NOVEL LOW-SHEAR STRESS BIOREACTOR

**DOI:** 10.64898/2026.05.06.723052

**Authors:** Madison Bergmann, Nathan Belliard, Patrice Meunier, Baptiste Roumezi, Olivier Detournay, Ali G Turhan, Annelise Bennaceur-Griscelli

## Abstract

**Background:** The use of autologous or allogeneic cell therapies has now entered to the clinical practice in several fields of medicine, especially in oncology and hematology. From this regard, 2D-cell manufacturing is complex and costly and bioreactors have attracted major interest for efficient and cost-effective mass production of cells.

Bioreactors have several advantages such as homogeneous repartition of nutrients and gas, control of all culture parameters and increased yield. However, the important shear stress generated by those bioreactors is an important disadvantage as it can affect cell survival or cell quality. This important shear stress is the result of the mixing method using either blades (used in stirred-tanked bioreactors) or gas bubbles (used in airlift bioreactors). Another downside of the use of bioreactors is the difficulty to scale-up. As the volume increases, the shear stress generated by blades radically increases leading to cell death and a decrease of cell quality.

**Description:** In this study, we describe a bioreactor developed using a different mixing method effectively reducing the shear stress and facilitating scale-up. This bladeless method uses an inclination of the bioreactor as well as rotation to mix fluids in a container. Here we described different steps that led to the adaptation of this bioreactor, initially developed for fragile microalgae culture, for mammalian cell culture amplification. The bioreactor was tested to amplify a natural killer (NK) cell line NK92 which is an IL-2 dependent cell line used in clinical trials for cancer therapy. We have tested the influence of 1-The number of cells seeded; 2-The influence of the rotation speed on cell growth and viability; 3-The influence of the bioreactor angle on the above parameters; 4-The duration of the culture.

**Results:** Cells were initially seeded at 2.5.10^5^ / ml in a volume of 380 ml. According to the rotation speed of 15, 30, 45 and 60 rpm, we have observed an increase of cell numbers at day 3 (3-fold), day 5 (7-fold) and day 7 (10-fold) compared to seeding, the best expansion being obtained at day 7 with a rotation speed of 45 rpm. The optimal angle of rotation was found to be 3 degree, with an optimal amplification at day 7 versus day 3 (p < 0.01). The viability was also found to be optimal in the latter condition.

**Conclusions:** These preliminary results demonstrate that NK92 cells could be amplified using this bioreactor. In the best tested condition, neither cell viability nor cell growth was impacted. These results strongly suggest the potential use of this device in future clinically applicable conditions.

## INTRODUCTION

In recent years, bioreactors have emerged as a novel strategy to efficiently accomplish a transition from laboratory-scale cell cultures to industrial-scale cell cultures in the field of cell therapy (1, 2, 3). In addition, in the growing industrial cell therapy field in cancer, there is major unmet need for efficient manufacturing and expansion of cells in cGMP grade. For example, the biopharmaceutical industry needs such cells for protein (monoclonal antibodies, hormones), vaccine and exosome production. In regenerative medicine, stem cells or differentiated cells are required to manufacture 3D organoids in cGMP conditions prior to potential transplantation. Organoids generated from tissue samples or from induced pluripotent stem cells (iPSCs) are currently used for disease modelling and drug screening but the 2D conditions do not allow the accurate conditions to mimic the organizational structure of a tissue. The successful generation of 3D organoids could be a major progress in this field allowing clinical developments.

Cultures in 2D conditions show low productivity and poor control of culture parameters. Moreover, important variation in cell quality can be observed due to manual handling. These drawbacks make traditional approaches unsuitable for large-scale production of clinical-grade cells due to inconsistent yield and quality and increased risks of contamination.

To accommodate the increasing needs in cells and avoid these drawbacks, bioreactors have been established as the standard solution. A variety of bioreactors have been developed to accommodate the specific needs of cells and achieve large scale production while maintaining cell quality. Currently, the most widespread bioreactors are the stirred tank bioreactors, using mechanical agitation by impellers to achieve gas, nutrient and cell homogenization. Despite the efficient control of the cellular environment, the use of impellers creates important shear stress that can be detrimental to cell viability, growth and expansion especially for fragile cells such as induced pluripotent stem cells (iPSCs).

Here, we report the adaptation of a bioreactor initially developed for microalgae (Dinoflagellates) culture.

This bioreactor developed at Planktovie is based on the SoftMixer principle described by Meunier ( 4 ) and Lefranc et al ( 5 ). In this system, mixing is achieved not by an internal impeller but by the rotation of the culture vessel around its axis, while the vessel is slightly inclined. This configuration generates a global movement of the liquid and suspended cell aggregates. To obtain the desired hydrodynamic behavior, the reactor must respect a specific aspect ratio (H/D, liquid height to vessel diameter), and the rotation speed must be carefully adjusted according to the working volume. Under these conditions, the system produces a chaotic turbulent-like flow regime that ensures efficient and homogeneous mixing while maintaining very low shear stress, which is particularly suitable for sensitive suspension cultures such as NK-92 cells. Oxygen transfer occurs mainly through gas exchange at the liquid surface, avoiding sparging and bubble formation and thereby preserving cell viability and functionality.

After adaptation to mammalian cells cultures using different cell lines, the bioreactor was tested to determine optimal culture condition for NK-92 cells. NK-92 cells have been a major alternative to autologous natural killer (NK) cells since their initial description in 1990 ( 6 ). This cell line, generated from the peripheral blood of a patient with aggressive lymphoma, has shown a major NK cell activity and has been since then in clinical trials after irradiation ( 7, 8) with several clinically applicable variants described ( 9, 10 ).

## MATERIALS AND METHODS

### Description of the bioreactor

Building on previous work on SoftMixer hydrodynamics (4, 5), we adapted this reactor concept for the culture of human cells. To do so, we used a cylindrical vessel with a diameter of 7.5 cm and a working volume of 380 mL, while maintaining an aspect ratio (H:D) of 1:1, as recommended for optimal mixing conditions in tilted rotating systems. Because the hydrodynamic regime is highly sensitive to the inclination angle of the vessel, several angles were experimentally evaluated (3°, 5°, and 10°, with a precision of 0.1°) in order to determine the configuration that allowed the fastest and most efficient resuspension of the cells while preserving homogeneous mixing conditions.

In addition, different rotation speeds (15, 30, 45, and 60 rpm) were tested to identify the operating conditions that provided optimal mixing while maintaining low shear stress suitable for human cell cultures (smaller than 2 mPa for the fastest speed ( 4 ).

A Trinamic Motion Control PD60-4-1276-TMCL stepper motor (200 steps subdivided into 256 microsteps) drives the bioreactor in rotation via a belt and toothed-wheel system with a reduction ratio of 60/15. This provides a high torque of 9.1 Nm, which compensates for the friction of the slip ring. Indeed, gases and electrical signals are supplied to the rotating bioreactor through a Mophlon slip ring MAPH200-01-P0610-S06. The gas consists of pure oxygen or pure CO₂, with injection controlled according to the pH and dissolved oxygen level of the culture medium, measured using an SFR Vario sensor (PreSens). The temperature of the bioreactor is maintained at 37 °C using a heating blanket manufactured by RS PRO.

### NK92 cell line

NK-92 cells were maintained in RPMI 1640 medium supplemented with 20% FBS, 1% penicillin/streptomycin and 50 U/mL of IL-2 (Miltenyi Biotec, #130-097-747) as previously described ( 11 ).

### NK92 Cell Culture in the Bioreactor

To make a comparison of the expansion potential of the bioreactor as compared to classical 2D conditions, NK-92 cells were cultured either in a 5% CO_2_ incubator in a flask with 20 ml of RPMI 1640 as described above or in the bioreactor with the adjusted volume of medium for 7 days. Gas injection parameters were determined experimentally to regulate pH and pO_2_. To maintain pH under 7.3, CO_2_ was injected up to every hour if needed with the schedule involving a 2 second injection if there is a 0.5 unit difference or more, and a 1second injection for a difference of 0.1 unit. To maintain pO_2_ above 85%, O_2_ was injected up to every 30 min if needed with the schedule involving a 0.2 second injection if the difference is 10% or more and a 0.1 second injection for a difference of 5%. Every other day, a half medium change was performed after cell counting, NK-92 cells were centrifuged and pelleted cells were returned to the culture.

### Proliferation assay

To determine growth and viability, comparative proliferation assays were performed in both static and dynamic (3D) conditions. For each test, two parameters (cell density, rotation speed and angle) were fixed and one was variable. Every other day, a half medium change was performed as described above. At each medium change, cells were counted using cell counter countess II FL (Invitrogen) and trypan blue for viability assessment (Invitrogen, #T10282). Mean value were expressed as cell concentration (cell/ml) and viable cell percentages (%).

### May Grünwald Giemsa staining

After collection of NK92 cells, 10^5^ cells were centrifuged on a slide using a cytofunnel (Thermo scientific, #5991040). Cells were then fixed and stained using cellavision RAL kit (#361550-0000) following suppliers instruction. Photographs were taken using an inverted microscope.

### Statistical analysis

ANOVA and paired student t-test were performed using GraphPad Prism version 10.0.0 for Windows, GraphPad Software, Boston, Massachusetts USA, www.graphpad.com. Means and SD are represented on the graphs. *, P < 0.05; **, P < 0.01; ***, P < 0.001.

## RESULTS

### 1 Adaptation of the Bioreactor for NK92 cell culture

The bioreactor for microalgae culture presented several technical issues that needed to be addressed prior to test its use in mammalian cell culture and amplification. Amongst these issues, the absence of temperature control, the injection of only one gas (CO_2_) and the difficulty to maintain a sterile culture environment were the most important points to modify.

The bioreactor used in microalgae cultures consisted of two elements including the automaton and the culture platform. The automaton underwent very few modifications as it was just reduced to accommodate injection of two gases from bottles without mixing them. The culture platform, however, underwent major modifications.

The adapted culture platform can also be divided in two elements: a static and a rotational element (Figure 1). The static element is constituted of a platform necessary to adjust the angle, a motor and a slip ring. The slip ring allows the passage of gas and liquid from the static to the rotary element.

**FIGURE 1.**
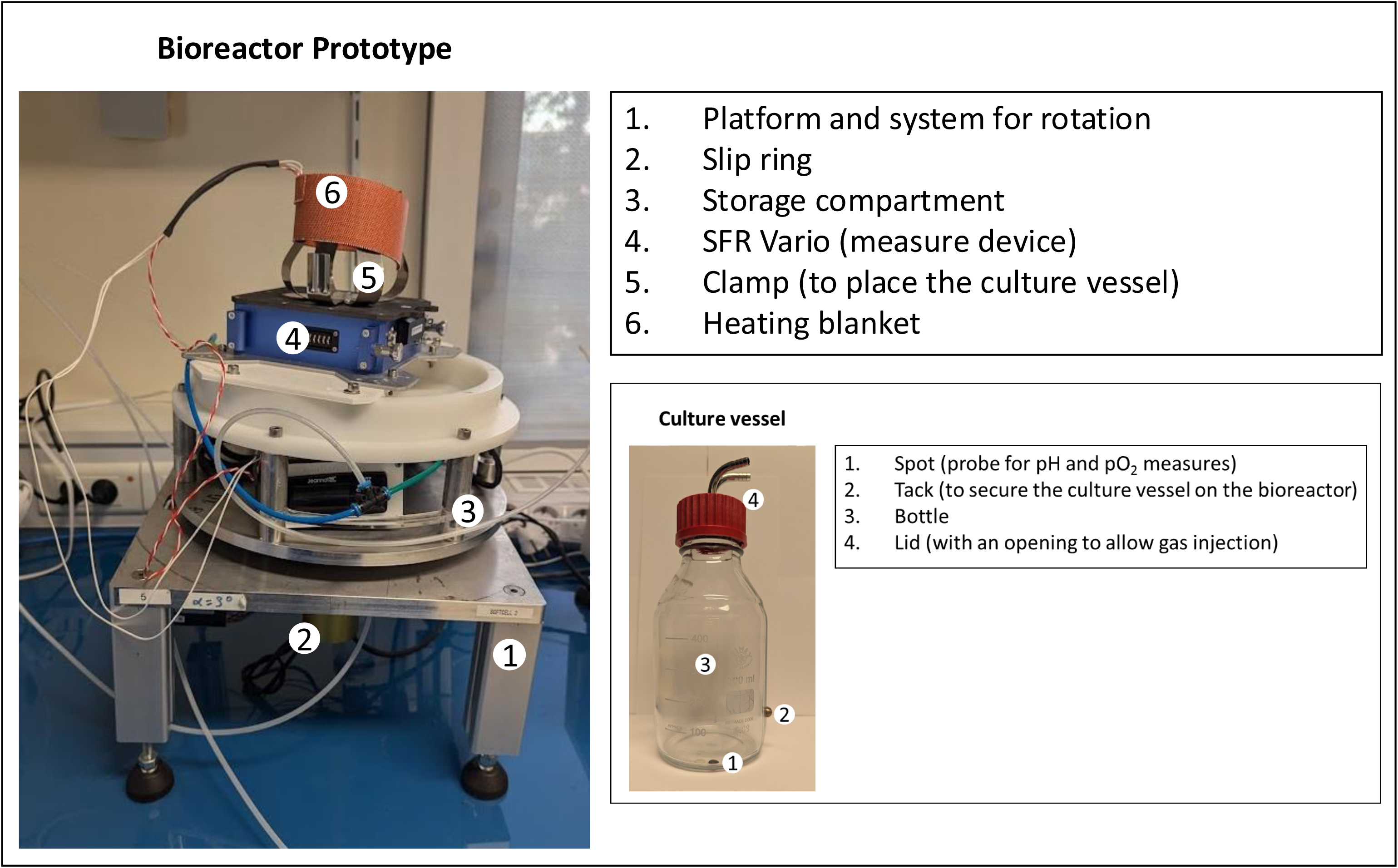
Photograph of the prototype of the Biorecator (on the left) and the culture vessel (on the right). Bottom part of the bioreactor (1 and 2) are static elements, top part of the bioreactor (3 to 6) are rotary elements. The culture vessel is placed on the clamp. The probes of the culture vessel should be aligned on top of the reading space of the SFR Vario.

The heating blanket was positioned in the storage compartment on top of which a measuring device was placed. This device (SFR Vario, System c bioprocess) allows real-time measurement of pH and oxygen in the culture media using non-invasive probes placed in the culture vessel. In the prototype version used, the culture vessel was a glass bottle that can be sterilized using an autoclave (Figure 1). The cap has two inset allowing gas injection. A 0.22µm filter was added at the junction between the inset and the pipe and the heating blanket was placed around the bottle to maintain the cell suspension at 37°C. Figure 1 shows the final prototype used in these studies.

Tests were performed to confirm the sterility of the culture as well as the maintenance of the temperature. Gas injection parameters were also tested and confirmed. Injection of CO2 occurred when needed and up to every hour to regulate pH and maintain it below 7.3 whereas injection of O_2_ took place every 30 minutes to regulate pO2 and keep it above 80%. Duration of each injection depended on the difference between the measurement and the set value.

### 2 Evaluation of physical parameters on cell growth

Figure 2 shows the design of the experimental strategy. After the initial cultures in flasks, the NK92 cells were divided in two equal fractions and were kept in culture for seven days for the first experimental set up used for growth curves.

**FIGURE 2.**
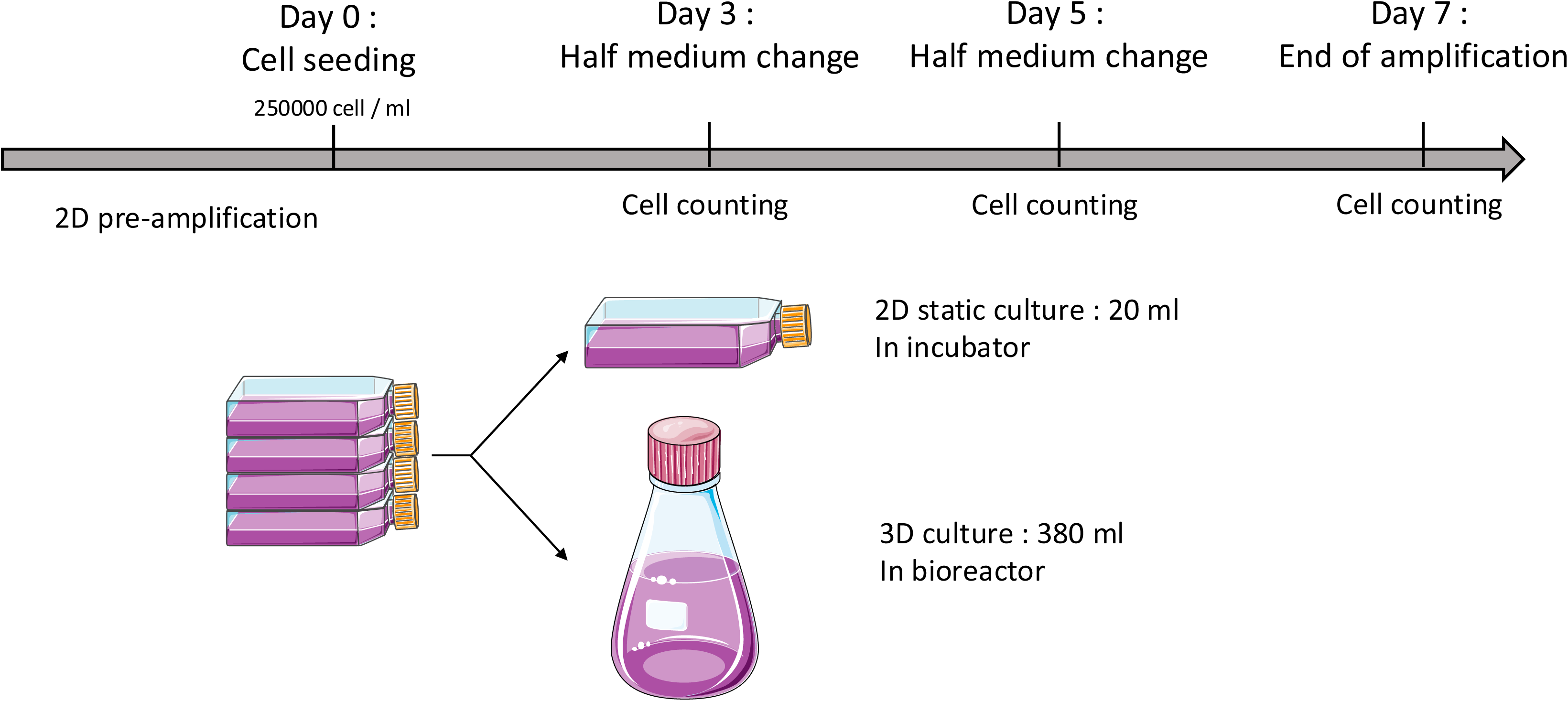
Design of experimental strategy (see methods)

#### a. Rotation speed

As previously described, the rotation speed is an important parameter that influences cell homogenization within the culture vessel. Four different speeds, ranging from 15 to 60 rpm, were evaluated on both cell growth and viability as compared to a static 2D control condition. Cells were seeded at 2.5.10^5^ cell / ml in the appropriate volume and the bioreactor was set at a fixed 3° angle. The volume was set at 20 mL for the 2D-control in optimal culture condition in an incubator and 380 ml for the bioreactor.

At the lowest speed tested (15 rpm) the cell suspension remained heterogenous with the majority of cells accumulating at the bottom of the culture vessel, indicating insufficient resuspension. In contrast, homogeneous cell suspensions were observed at all higher rotation speeds tested.. This heterogeneity at 15 rpm was associated with a marked reduction in cell proliferation, resulting in only a 4-fold amplification by day 7 (figure 3A), compared to a 9.5-fold expansion in the control static 2D control condition. Using this low-speed condition, cell viability was also significantly reduced, reaching 91.5 ± 2.69 % at day 5 and 89.8 ± 2.47 % at day 7, whereas viability remained above 95 % under all other tested conditions (figure 3B).

**FIGURE 3.**
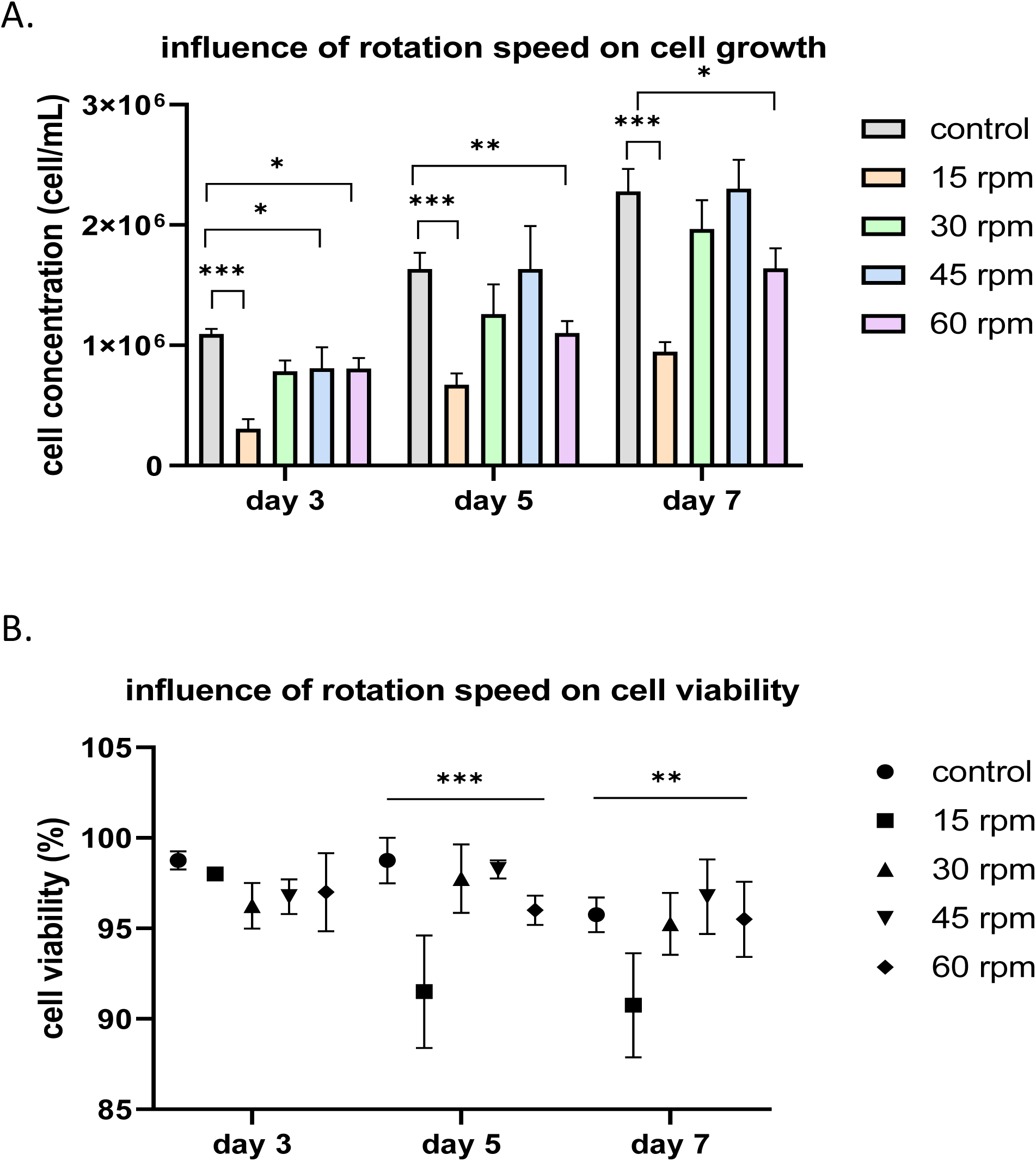
Evaluation of the cell growth and viability of NK-92 cells according to different rotation speeds with a fixed angle of 3 degrees as compared to control conditions (Gray columns, 2D static condition). A. Influence of rotation speed on cell growth B. Influence of rotation speed and cell viability. * p < 0.05, ** p < 0.01, *** p < 0.001

At the highest speed tested (60 rpm), cell proliferation was slightly reduced, with a 7-fold expansion at day 7 (figure 3A) compared to a 9.5-fold expansion in the static 2D control (condition static 2D). However, no effect was observed on cell viability (figure 3B).

Two intermediate rotation speeds were also tested. At 30 rpm, cells exhibited a 9-fold expansion, while at 45 rpm the expansion reached 9.9-fold by day 7. In both cases, cell growth was comparable to the control condition (9.5-fold expansion ) although the volume is 20 times larger in the bioreactor than in the control flask.. The highest proliferation rate was observed at 45 rpm which yielded the best overall expansion among the tested rotation speeds..

#### b. Bioreactor angle

The inclination angle represents a second key physical parameter influencing homogenization of the cell suspension, and its impact on cell culture performance was therefore evaluated. Increasing the angle allowed a better homogenization at low speed without increasing shear stress. To test the effect of the inclination on cell growth, three angles (3, 5 and 10 degrees) were tested at a 30 rpm rotation speed and a fixed 2.5.10^5^ cell/ml seeding.

At a 3 degree inclination, cell expansion was comparable to that observed in the control condition (figure 4A), without any effect on cell viability (figure 4B).

**FIGURE 4.**
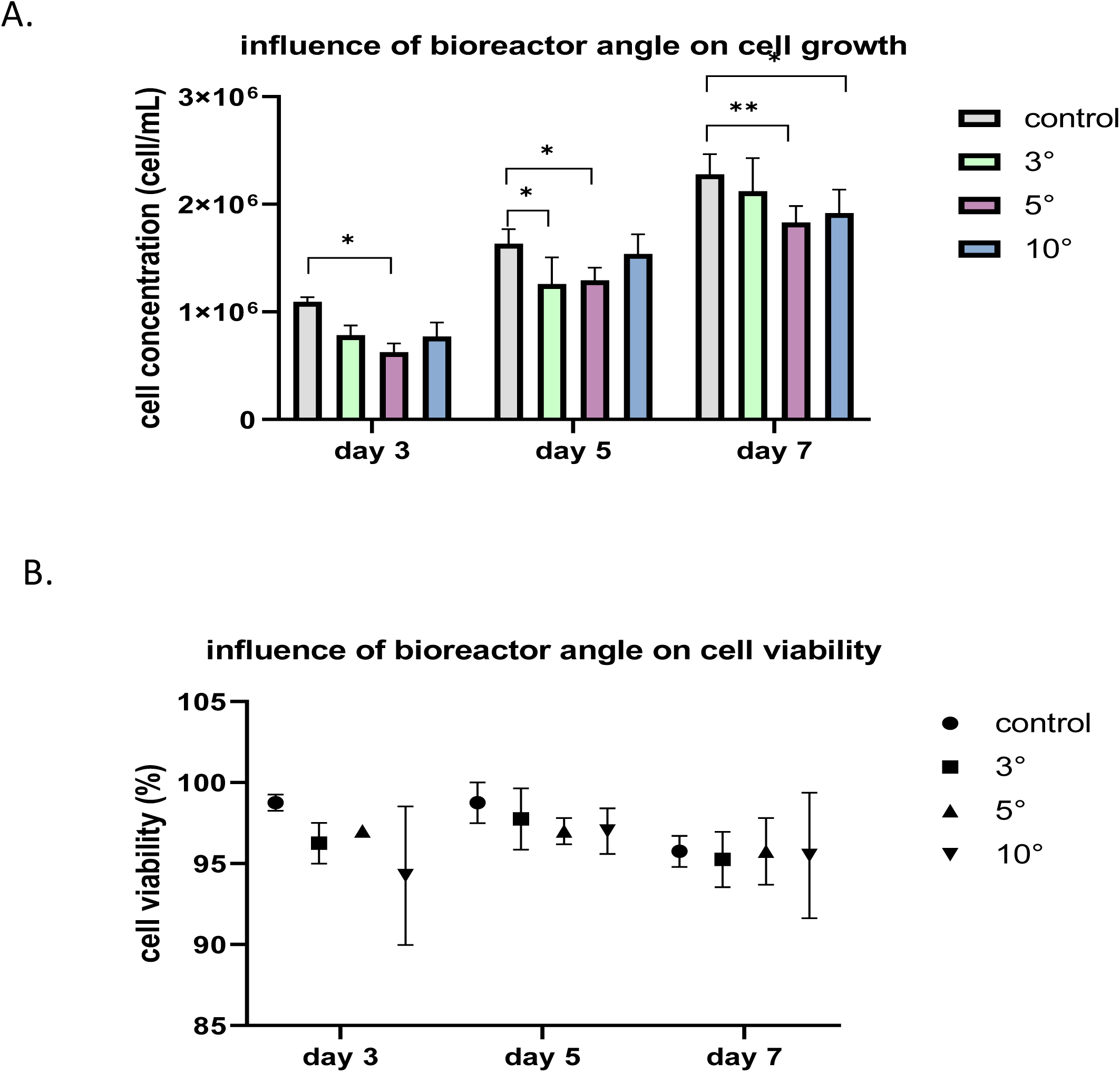
Evaluation of the growth of NK92 cells according to the bioreactor angle with affixed rotation speed of 30 rpm as compared to static (2D) condition (Gray columns ). A. Influence of the angle of rotation on cell growth. B. Influence of the angle of rotation on cell viability at different angles (fixed rotation speed of 30 rpm) as compared to control (2D static condition). * p < 0.05, ** p < 0.01, *** p < 0.001

Increasing the angle at 5 degrees or 10 degrees had no effect on cell viability (figure 4B), which remained above 95% as found in the control condition. However, an effect on cell growth can be observed at day 7 for both angles (figure 4A). At an angle of 5 degrees, a 8.3-fold amplification was found and a 8.4-fold amplification was found at a 10 degrees as compared to the 9.5-fold amplification found in the control condition.

### 3 Morphological characteristics of the cells post-expansion

Under the optimal culture conditions identified ( 45 rpm rotation at a 3 degree angle), macroscopic observation shows the presence of aggregates in the bioreactor. This observation is confirmed microscopically. In control 2D culture condition, few aggregates can be observed and their size is heterogenous. However, in the bioreactor, many smaller aggregates can be observed (figure 5 A). May Grunwald Giemsa staining revealed no noticeable differences in cells after 3 days in culture in either 2D static culture condition or 3D culture condition (figure 5B).

**FIGURE 5.**
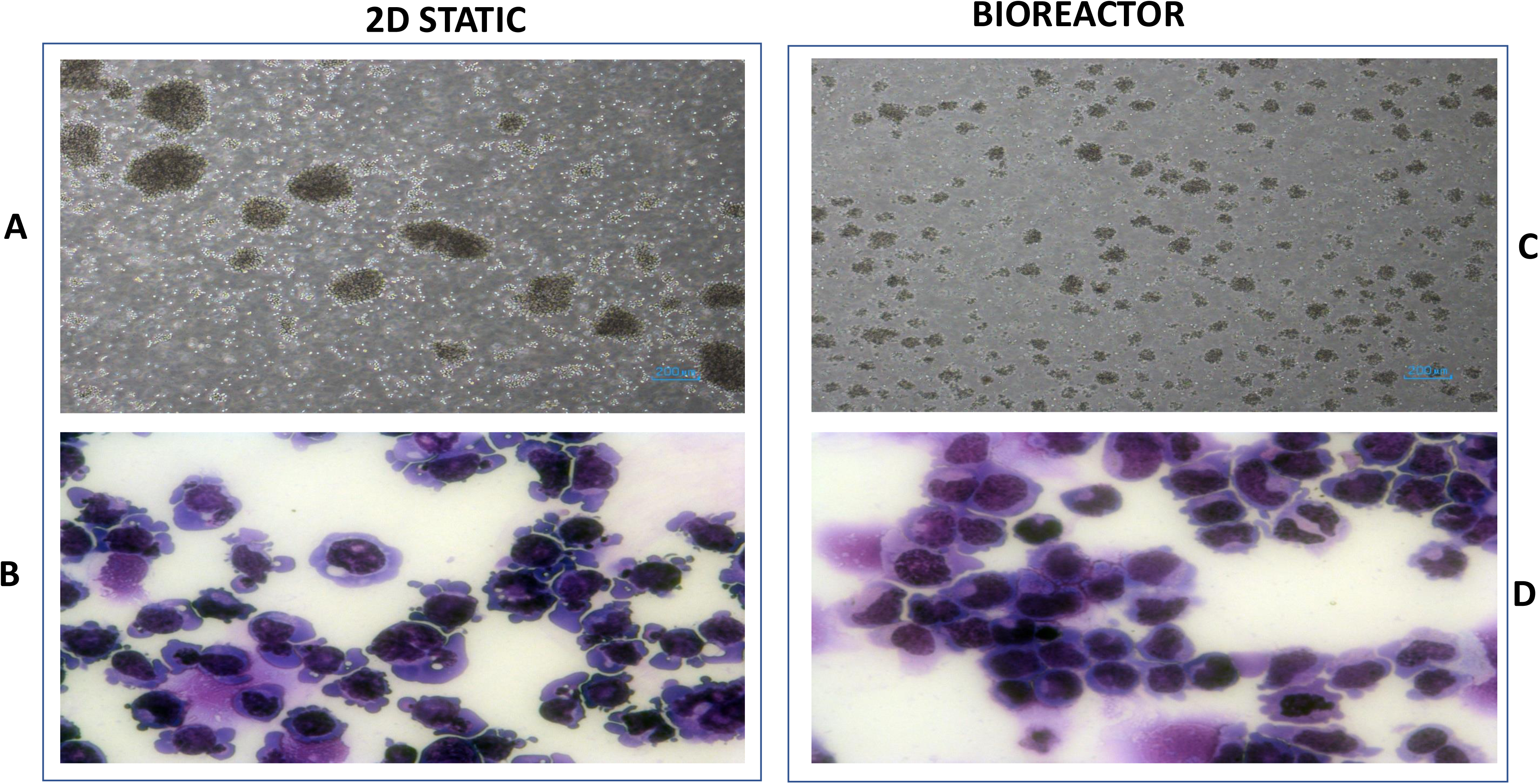
Microscopic observation of NK92 cells at day + 3 days of culture in the control 2D flask (Left panel) or in the bioreactor (Right panel). A. Contrast microscopy in 2D Flasks B. May-Grünwald Giemsa staining in 2D C. Contrast microscopy of cells grown in the Bioreactor D. May-Grünwald Giemsa staining of cells grown in the Bioreactor.

## DISCUSSION

We report here the characteristics of a 3D bioreactor which has been used for the first time to expand the NK92 cell line which is a major immunotherapy tool used in clinical trials ( 8 ). Studies aiming to amplify NK92 cells using bioreactor technologies have been previously published (12, 13, 14). In the study published by Lin et al, static conditions were compared to a mechanical rotation (15 rpm) for several hours before returning the cells to static conditions. The cell aggregate size was found to be reduced and the viability was found to be increased ( 12 ). In the work published by Lee et al a closed bioreactor system by Cytiva was used to test the expansion and viability after cryopreservation. The expansion capacity after thawing was found to be increased using the bioreactor system but the flask-mediated expansion remained easier to use and less costly, making it a practical and efficient option for routine low scale cell expansion ( 13 ).

The bioreactor described in this work compares favorably with conventional bioreactors systems such as stirred-tank, wave or airlift bioreactors which can generate significant shear stress as a result of mechanical agitation, aeration or medium circulation. Shear stress is known to influence cell physiology and quality and, in sensitive cell type it may even trigger apoptosis. For this reason, tight control of hydrodynamic stress is a critical parameter in large-scale cell expansion. To mitigate these effects, carrier-and bead-based culture systems have been developed. In these systems, cells are placed either at the surface or in a scaffold that protects them from shear stress while allowing the use of a bioreactor for cell amplification. While these systems are proven to maintain cell viability and quality, their use reduces the benefits of the bioreactor as they can limit nutrient and gas exchange and reduce the volume of medium available for amplification.

The key advantage of the monitoring of these parameters is the fact that they can be used to reflect on the culture in real time. It was seen experimentally that O_2_ consumption in the culture correlates with cell growth and viability. As cells divide, O_2_ intake increases and more injections are needed to maintain 80% oxygen in the medium. However, when cells are not dividing or are undergoing apoptosis, this uptake decreases and less injections are needed. The same phenomenon can be seen with pH measurement. As cells grow, nutrients in the medium are used and pH decreases. Taken together these data suggest that pH and pO_2_ measurement is an reliable real time monitoring of cell health in culture.

Another advantage to the modification of culture vessel and finer gas control was the effect on medium evaporation. As the gas injections are reduced in number and duration, the medium remains in the culture vessel.

Overall, the adapted bioreactor (figure 1) underwent major modifications to make it suitable for human cell culture applications. The final design operates at a working volume of 380 ml which is 20 times larger than the control static flask. One of the main advantages of this system lies in its intrinsic scalability. In the field of cell therapy manufacturing, the development of culture systems allowing efficient scale-up is essential to reduce production costs and to ensure greater batch-to-batch consistency, both of which remain major challenges for the industrialization of cell-based therapies ( 15 )

In conventional bioreactor systems, increasing the working volume often leads to higher hydrodynamic stresses, due to stronger agitation, aeration or medium circulation, which can complicate scale-up and negatively affect sensitive cells. In contrast, the bioreactor described here relies on the SoftMixer hydrodynamic principle, for which sear stress remains constant as the culture volume increases (4, 10). This counterintuitive but advantageous property makes the system particularly well suited for progressive scale-up of cell cultures, while maintaining low mechanical stress and homogeneous culture conditions.

Two physical parameters can influence cell growth and viability: speed of rotation and inclination of the bioreactor. Both parameters were tested to evaluate their effect on cell culture and determine the optimal culture conditions.

Speed of rotation had the largest influence on both cell growth and viability. At low speed, an heterogenous suspension was found which resulted in cell death probably due to apoptosis. In contrast, at higher rotation speeds, cell proliferation in the bioreactor was very similar to the static case. However, the growth rate was slightly reduced for the highest speed (by 25%) or the highest inclination angle (by 10%). NK-92 cells are known to naturally grow in aggregates and to depend on cell–cell interactions for optimal survival and proliferation, while they can undergo cell death when excessively dispersed (16 ). Excessive mixing may therefore limit the formation of these transient aggregates, resulting in reduced cell amplification. Consistent with this hypothesis, small cellular aggregates were observed during the first days of culture at 30 and 45 rpm, suggesting that these intermediate hydrodynamic conditions favored cell–cell contacts. However, by day 7, aggregates were no longer observed in either the control or the bioreactor culture conditions, indicating that aggregation was transient during the expansion phase.

We have found that increasing the angle led to a decrease in cell amplification. When compared to the amplification observed at 60 rpm, the stunt cell growth was not found to be important potentially related to the use of a higher angle inclination, leading to the formation of aggregates.

We also show the optimal culture conditions as tested are 3° angle and 45 rpm rotation. Other cell culture related parameters should be tested such as cell density seeding and IL-2 concentration in the medium. As it was reported that NK-92 cells may require more IL-2 for their growth in 3D culture condition.

S everal studies reported that cells often require a period of adaptation when transferred from static culture systems to bioreactor environments (ref).. In the present study, experiments were conducted over 7 days, and each run was initiated using cells freshly expanded under 2D culture conditions. It is therefore possible that longer-term culture or successive passages in the bioreactor could further improve cell expansion once cells become adapted to the hydrodynamic conditions of the system.

Beyond the continued clinical interest of NK cells and their derivatives (15, 16 ) the use of 3D bioreactors is rapidly expanding for the large scale culture of induced pluripotent stem cells ( 17 ) and organoids ( 18, 19, 20). With regard to the expansion of fragile cells such as iPSCs, the shear stress is a major limiting factor. Indeed, several recent works are aiming to reduce this event with the goal of improving expansion of iPSCs ( 21, 22) In this context, our results demonstrate for the first time that a low-shear-stress bioreactor based on the SoftMixer principle represents a promising approach for scalable cell culture. This bioreactor creates a shear stress which is 10 to 100 times smaller than in a classical stirred-tank bioreactor (4). Such a system could be particularly advantageous not only for the expansion of NK cells, but also for the culture of iPSCs and their derivatives, thereby supporting the development of next-generation cell therapies currently being explored in oncology and regenerative medicine. Importantly, improving the scalability and robustness of cell manufacturing processes is essential to reduce production costs, which currently represent one of the major barriers limiting the widespread clinical adoption of cell therapies and preventing a larger number of patients from benefiting from these treatments..

## ACKNOWLEDGMENTS

This work has been supported by the grant **N° DOS0166813/00** “Grand Defi Bioproduction” France 2030 from French Ministry of Research, ANR and BPI and additional Grants and support from INSERM, Paris Saclay University and UMRS 1310.

## REFERENCES

1. Baghbaderani BA. Commercialization of Cell Therapies: A CDMO Perspective. Adv Exp Med Biol. 2025;1486:137–145. doi: 10.1007/978-3-031-97297-3_11. PMID: 41136837.

2. Silva Couto P, Stibbs DJ, Springuel P, Schultz U, Effenberger M, Goldrick S, Navarro-Velázquez S, Juan M, Herbst L, Nießing B, Mestermann K, Sanges C, Hudecek M, Rafiq QA. Impact of Serum/Xeno-Free Medium and Cytokine Supplementation on CAR-T Cell Therapy Manufacturing in Stirred Tank Bioreactors. Biotechnol J. 2025 Sep;20(9):e70114. doi: 10.1002/biot.70114. PMID: 40923846; PMCID: PMC12419138.

3. Scibona E, Morbidelli M. Expansion processes for cell-based therapies. Biotechnol Adv. 2019 Dec;37(8):107455. doi: 10.1016/j.biotechadv.2019.107455. Epub 2019 Oct 17. PMID: 31629791.

4. Meunier P. (2020) Geoinspired soft mixers. Journal of Fluid Mechanics. 2020;903:A15. doi:10.1017/jfm.2020.634

5. Lefranc C, Detournay O, Meunier P ; Gas injection into a tilted rotating cylinder. Physics of Fluids 1 June 2023; 35 (6): 066602

6. Gong J-H, Maki G, Klingemann HG. Characterization of a human cell line (NK-92) with phenotypical and functional characteristics of activated natural killer cells. Leukemia. 1994;8(4):652–8.

7. Arai S, Meagher R, Swearingen M, Myint H, Rich E, Martinson J, Klingemann H. Infusion of the allogeneic cell line NK-92 in patients with advanced renal cell cancer or melanoma: a phase I trial. Cytotherapy. 2008;10(6):625–32.

8. Tonn T, Schwabe D, Klingemann HG, Becker S, Esser R, Koehl U, Suttorp M, Seifried E, Ottmann OG, Bug G. Treatment of patients with advanced cancer with the natural killer cell line NK-92. Cytotherapy. 2013;15(12):1563–70.

9. Klingemann H. The NK-92 cell line-30 years later: its impact on natural killer cell research and treatment of cancer. Cytotherapy. 2023 May;25(5):451–457. doi: 10.1016/j.jcyt.2022.12.003. Epub 2023 Jan 6. PMID: 36610812.

10. Klingemann H. The Natural Killer Cell Line NK-92 and Its Genetic Variants: Impact on NK Cell Research and Cancer Immunotherapy. Cancers (Basel). 2025 Jun 13;17(12):1968. doi: 10.3390/cancers17121968. PMID: 40563618; PMCID: PMC12191011.

11. Imeri J, Marcoux P, Huyghe M, Desterke C, Fantacini DMC, Griscelli F, Covas DT, de Souza LEB, Griscelli AB, Turhan AG. Chimeric antigen-receptor (CAR) engineered natural killer cells in a chronic myeloid leukemia (CML) blast crisis model. Front Immunol. 2024 Jan 8;14:1309010. doi: 10.3389/fimmu.2023.1309010. PMID: 38259442; PMCID: PMC10801069.

12. Lin JN, Kuan CY, Chang CT, Chen ZY, Kuo WT, Lin J, Lin YY, Yang IH, Lin FH. High-throughput proliferation and activation of NK-92MI cell spheroids via a homemade one-step closed bioreactor in pseudostatic cultures for immunocellular therapy. J Biol Eng. 2024 Nov 12;18(1):65. doi: 10.1186/s13036-024-00461-0. PMID: 39533411; PMCID: PMC11555828.

13. Lee S, Joo Y, Lee EJ, Byeon Y, Kim JH, Pyo KH, Kim YS, Lim SM, Kilbride P, Iyer RK, Li M, French MC, Lee JY, Kang J, Byun H, Cho BC. Successful expansion and cryopreservation of human natural killer cell line NK-92 for clinical manufacturing. PLoS One. 2024 Feb 23;19(2):e0294857. doi: 10.1371/journal.pone.0294857. PMID: 38394177; PMCID: PMC10889882.

14. Huang H, Zhang S, Zhao Y, Xu R, Tan WS, Cai H. Suspension culture promoted the expansion of NK-92 cells ex vivo by enhancing the expression of IL-2 receptor. Biotechnol J. 2024 Mar;19(3):e2300654. doi: 10.1002/biot.202300654. PMID: 38472089

15. Cherian DS, Bhuvan T, Meagher L, Heng TSP. Biological Considerations in Scaling Up Therapeutic Cell Manufacturing. Front Pharmacol. 2020 May 13;11:654. doi: 10.3389/fphar.2020.00654. PMID: 32528277; PMCID: PMC7247829.

16. Zhang J, Zheng H, Diao Y. Natural Killer Cells and Current Applications of Chimeric Antigen Receptor-Modified NK-92 Cells in Tumor Immunotherapy. Int J Mol Sci. 2019 Jan 14;20(2):317. doi: 10.3390/ijms20020317. PMID: 30646574; PMCID: PMC6358726.

17. Francis N, Aho J, Ben-Nun IF, Bharti K, Dianat N, Makovoz B, Nouri P, Rothberg J, Song H, Zamilpa R, Lakshmipathy U, Allickson J. Scaling up pluripotent stem cell-based therapies - considerations, current challenges and emerging technologies: perspectives from the ISCT Emerging Regenerative Medicine Working Group. Cytotherapy. 2025 Sep;27(9):1031–1042. doi: 10.1016/j.jcyt.2025.04.058. Epub 2025 Apr 7. PMID: 40353785.

18. Vicente P, Inocêncio LR, Ullate-Agote A, Louro AF, Jacinto J, Gamelas B, Iglesias-García O, Martin-Uriz PS, Aguirre-Ruiz P, Ríos-Muñoz GR, Fernández-Santos ME, van Mil A, Sluijter JPG, Prósper F, Vega MMM, Alves PM, Serra M. Billion-Scale Expansion of Functional hiPSC-Derived Cardiomyocytes in Bioreactors Through Oxygen Control and Continuous Wnt Activation. Adv Sci(Weinh). 2025 Mar;12(11):e2410510. doi: 10.1002/advs.202410510. Epub 2025 Jan23. PMID: 39846380; PMCID: PMC11923921.

19. Lecina M, Ting S, Choo A, Reuveny S, Oh S. Scalable platform for human embryonic stem cell differentiation to cardiomyocytes in suspended microcarrier cultures. Tissue Eng Part C Methods. 2010 Dec;16(6):1609–19. doi: 10.1089/ten.TEC.2010.0104. Epub 2010 Aug 28. PMID: 20590381.

20. Teo A, Mantalaris A, Song K, Lim M. A novel perfused rotary bioreactor for cardiomyogenesis of embryonic stem cells. Biotechnol Lett. 2014 May;36(5):947–60. doi: 10.1007/s10529-014-1456-y. Epub 2014 Mar 21. PMID: 24652542.

21. Kim J, Agbojo O, Jung S, Croughan M. Measurement of Oxygen Transfer Rate and Specific Oxygen Uptake Rate of h-iPSC Aggregates in Vertical Wheel Bioreactors to Predict Maximum Cell Density Before Oxygen Limitation. Bioengineering (Basel). 2025 Mar 22;12(4):332. doi: 10.3390/bioengineering12040332. PMID: 40281691; PMCID: PMC12024368.

22. Teale MA, Schneider SL, Seidel S, Krasenbrink J, Poggel M, Eibl D, Sousa MFQ, Eibl R. Expansion of induced pluripotent stem cells under consideration of bioengineering aspects: part 2. Appl Microbiol Biotechnol. 2025 Feb 6;109(1):38.doi: 10.1007/s00253-024-13373-2. PMID: 39912924; PMCID: PMC11802622.

